# Analysis of the abomasal transcriptome of LDA affected cattle

**DOI:** 10.1101/2022.09.27.509652

**Authors:** Zoltán Gál, Bálint Biró, Zsófia Nagy, Levente Kontra, András Horváth, Orsolya Ivett Hoffmann

## Abstract

Left displacement of the abomasum (LDA) is a common condition in Holstein population mainly occur around the time of parturition. The entrapped abomasum located between the rumen and the abdominal wall caused by the abomasal hypomotility. The heritability of LDA estimated higher than for other bovine diseases but a number of management and nutritional conditions are also affecting the disease appearance. Genome studies revealed many significant genomic regions associated with LDA, although an RNA sequencing analysis of abomasum is missing from the literature. Within the framework of this research, we tried to patch up this area missing from the literature and to reveal the genetic causes and a complex interaction between the endocrine and neuromuscular pathways behind the symptoms of the disease with the help of transcriptomic analysis.

## Introduction

Displacement of the abomasum is a frequent condition in dairy cattle which occurs due to abomasal hypomotility and gas accumulation. Most of the cases the abomasum replaced to the left side, although right displaced abomasum (RDA) is also existed (Hamann et al., 2004; Zerbin, Lehner & Distl, 2015). Left displaced abomasum (LDA) is a common dairy cattle disease occurring in 1-7% of the Holstein-Friesian population and in many other breeds. The LDA is commonly associated with high yielding and intensively fed dairy cows in early or late lactation, but the disease can be found in beef cattle populations as well (Hamann et al., 2004; Mömke et al., 2013). Effective prediction would be necessary as LDA increases veterinary costs of dairy farms and decreases life expectancy with restricted milk production (Mömke et al., 2012).

The multifactorial cause of the LDA presumably based on genetic factors affecting the initiation and maintenance of the MMC. The heritability estimates mostly measured in Holsteins fluctuate between 0.2 and 0.5 (Mömke et al., 2012; Mömke et al., 2013; Zerbin, Lehner, & Distl, 2015). LDA can occur in any time of the lifespan, however 80% of the displacements are observed within one month of partition (Wolf et al., 2001). Reduced abdominal motility is the primer etiological cause of the LDA and some breeding technology (like calving problems, overfeeding in the dry period, decreased feed intake) risk factors strongly affecting the incidence of the disease. Hypocalcaemia, metabolic alkalosis and concurrent diseases such as ketosis and metritis may also led to the development of the disease (Van Winden, S. C., & Kuiper, R., 2003). The occurrence of LDA may be reduced, but not completely eliminated by the avoidance of the risk factors. Some risk factors have been identified to play role in the evolving of the displacement of the abomasum like calving problems, twin births, overfeeding in the dry period, decreased feed intake and high body condition before calving (Van Winden, S. C., & Kuiper, R., 2003).

The migrating motor complex (MMC) is considered as a cyclic interdigestive reflex which traveling along the gastrointestinal track to propel and emptying those contents. The MMC comprehend three or four phases vary in species. Phase III is the most characteristic one contains intense contractions, the whole MMC cycle ranges between approximately 90-120 min. Among other peptides - like ghrelin, gastrin, serotonin - motilin is considered as an endocrine regulator of the MMC (Poitras & Peeters, 2008). In 1966 Brown et al. published a paper in which they described that the activity of denervated and transplanted pouches was elevated after the administration of alkaline solution. Two alternative explanations have emerged to interpret this phenomenon. One was that the alkaline solution contributes to the secretion of a stimulatory humoral agent. In those days they were not able to name this so-called humoral agent (Brown et al., 1966). After some years the same research group has successfully isolated this agent and they named it motilin after the observation that this polypeptide stimulates motor activity (Brown et al., 1971). Motilin is a small, 22 amino-acid peptide hormone encoded by the motilin (MLN) gene located at bovine chr23. This hormone plays an important role in the regulatory system of the interdigestive motility. It is produced by the endocrine cells of duodeno-jejunal mucosa and its plasma levels increase during the interdigestive periods. The peaks in the motilin plasma level leads to cyclical peristaltic contractions in the fasting periods in order to gastric emptying of the gut content (phase III of MMC). It is clear that there is a strong correlation between the expression patterns of motilin and the emergence of LDA. Interestingly a single nucleotide polymorphism (SNP) was found in the first non-coding exon of the bovine motilin gene. This SNP is now considered as a crucial influencer of the relation between the MLN gene and a transcription factor called NKX2-5. By the result of this mutation the MLN expression will be decreased by 89% compared to the wild type individuals. This SNP and the alteration of MLN expression may play an important role in bovine LDA through the motility of the abomasum (Mömke et al., 2012).

Nowadays it seems to be proven that gastrointestinal hormones such as ghrelin (GHRL), gastrin (GAST) and cholecystokinin (CCK) play a central role in the complex neuroendocrine interactions that became the basic principle in gastrointestinal motility regulation (Gué and Buéno, 1996).

GHRL is made up of 28-amino acids and it is synthesized in the upper gastrointestinal tract. It seems that ghrl has a function in the regulation of gastrointestinal motility (Ohno et al., 2010). Even though ghrl and motilin share high sequence similarity, ghrelin does not interact with the motilin receptor. What is more, the administration of ghrelin does not alter the plasma level of motilin (Deloose et al., 2019).

Gastrin (GAS) is a peptide hormone secreted by the G cells of pyloric antrum and duodenum (Romanski, 2017). It has a wide variety of biological functions for example in stimulating gastrointestinal motility, gastrointestinal mucosal growth and delaying gastric emptying (Misiewicz et al., 1969).

Cholecystokinin (CCK) is a polypeptide which shares a same C-terminal sequence with gastrin. This end of the peptides is responsible for the biological activity (Attoub et al., 1999). Cholecystokinin is produced by the I cells of the intestine. This hormone has several roles in biological processes including gallbladder, gastric emptying, pancreatic secretion and inhibition of food intake (Ivy and Oldberg, 1928; Deloose et al 2019). Interestingly, motilin level was not altered after intravenous cholecystokinin administration (Deloose et al., 2019).

Somatostatin (SST) is a neuropeptide hormone which has two sources. On one hand it is produced by the δ cells of Langerhans islets, and on the other hand it is produced by extraislet neuroendocrine cells (Hauge-Evans et al., 2009). It is involved in numerous processes including control of hormone secretion and inhibition of intestinal motility (Keskin & Yalcin, 2013). Activity of motilin release decreased after intravenous administration of somatostatin and its analogue both in human and dog (Deloose et al., 2019).

Pancreatic polypeptide hormone consists of 36 amino acids and it is produced by pancreatic islets. The two main biological effects of pancreatic polypeptide are the inhibition of the pancreatic exocrine secretion and the relaxation of the gallbladder (Leiter et al., 1984). Plasma level of motilin was decreased after pancreatic polypeptide administration in both dogs and humans (Deloose et al., 2019).

The essential amino acid tryptophan can be transformed through a complex pathway into 5-Hydroxytryptamine, better known as serotonin. This hormone is a monoamine neurotransmitter. The serotonergic neurons of the central nervous system and the enterochromaffin cells of the gastrointestinal tract are responsible for the production of serotonin. The enterochromaffin cells of the gastrointestinal tract contain more than 95% of the peripheral serotonin. Its role in the activation of neural reflexes and gastrointestinal tract peristalsis is crucial (Jonnakuty & Gragnoli, 2008).

Neurotensin is a peptide hormone containing 13 amino acids (Egloff et al., 2014). It is synthesized in the brain, in some peripheral organs and in the gastrointestinal cells. Absorptive processes were promoted after neurotensin release through several mechanisms containing increased ileal blood flow, pancreatic amylase and hepatic bile secretion. Furthermore, neurotensin regulates the motility of gastrointestinal tract by slowing down the transit time in the stomach and in the small intestine (Ratner et al., 2016).

There are just a few research about the expression patterns of different gastrointestinal hormones in ruminants. These experiments were mainly conducted on sheep. Gastrin expression was found to be the highest in the antral abomasum. High gastrin expression patterns were also found in the proximal and mid duodenum respectively (Reynolds et al., 1984). Ghrelin had the highest expression value in the abomasum (Huang et al., 2006). This result was partially confirmed by Wang et al. (2014) since they found that omasum and abomasum had the highest ghrelin expression values. Like the previous hormones, somatostatin also reached high expression levels in the abomasum. Beside abomasum, duodenal and pancreatic expression patterns were also considerable in terms of somatostatin (Darvodelsky et al., 1988).

Dual oxidase (DUOX) family has seven members in mammals, DUOX1-2, encoded by separate genes (Buvelot *et al*., 2019). The function of those enzymes is producing reactive oxygen species (ROS), superoxide anion, hydrogen peroxide (H2O2) and hydroxyl radicals in various tissues. Although, the seven members of the DUOX family share their function in ROS production, each has its own specific property. DUOX1, DUOX2 primarily generate H2O2. The sustained level of oxidative stress can lead to significant cell damage causing apoptosis or inflammation and play a significant role in the pathogenesis of chronic inflammation, activation of wound healing and tissue fibrogenesis. Since the oxidative stress is an imbalance between ROS generation and antioxidant defence, DUOX family plays prominent role in the regulation of cellular pathophysiology. Both the up- and downregulation of NOXs and DUOXs may contribute to the development of numerous diseases (Sirokmany *et al*., 2016; Zhang *et al*., 2020).

DUOX1 and 2 (previously called ThOX1 and ThOX2, respectively) are not expressed in the cardiovascular system in mammals (De Deken *et al*., 2000). DUOXs were identified in the epithelial cells in the thyroid gland, lower gastrointestinal (GI) tracts, respiratory system (Krause, 2004; Rada and Leto, 2008) and in the urothelium (Donko *et al*., 2010). DUOXA1 and DUOXA2 maturation factors are required for the endoplasmic reticulum to Golgi transfer and translocation to the plasma membrane of DUOX1 and DUOX2, respectively (Grasberger and Refetoff, 2006; Morand *et al*., 2009). Both DUOX1 and DUOX2 produce H2O2 in the thyroid gland, which is essential for proper iodination and thyroid hormone synthesis. Mutations in the DUOX genes or in their respective DUOXA partners, mainly in DUOX2/DUOXA2 genes, cause congenital hypothyroidism (CH), both in adult patients (De Deken and Miot, 2019; Weber *et al*., 2013) and children (Vigone *et al*., 2015; Wang *et al*., 2021).

Within the framework of this research, we examined which genes are expressed in the tissue of the bovine abomasum and compared the transcriptome of LDA-affected and healthy individuals. This study will complement the data of the expressing genes in the abomasum and therefore help us for the better understanding of the bovine genome and led to known better the genes involved in the development of the LDA disease.

## Results

To identify the factors affecting LDA, we used single end sequencing from the abomasum of 2 healthy and 5 LDA affected animals. Approximately 20 to 40 million reads were obtained for each library. The reads were aligned into the bovine UMD3.1 (GCA_000003055.3) reference genome.

PCA reduces the dimensions of the input matrix to define hidden patterns that can be used to cluster the input. PCA was performed to determine whether the control and LDA affected individuals can be clustered based on their gene expression profiles. Figure 1. shows that the principal component 1 (PC1) accounts for 55% of the total variation and hence control animals and LDA affected animals are grouped together based on their transcriptomic fingerprints.

**Figure 1.**
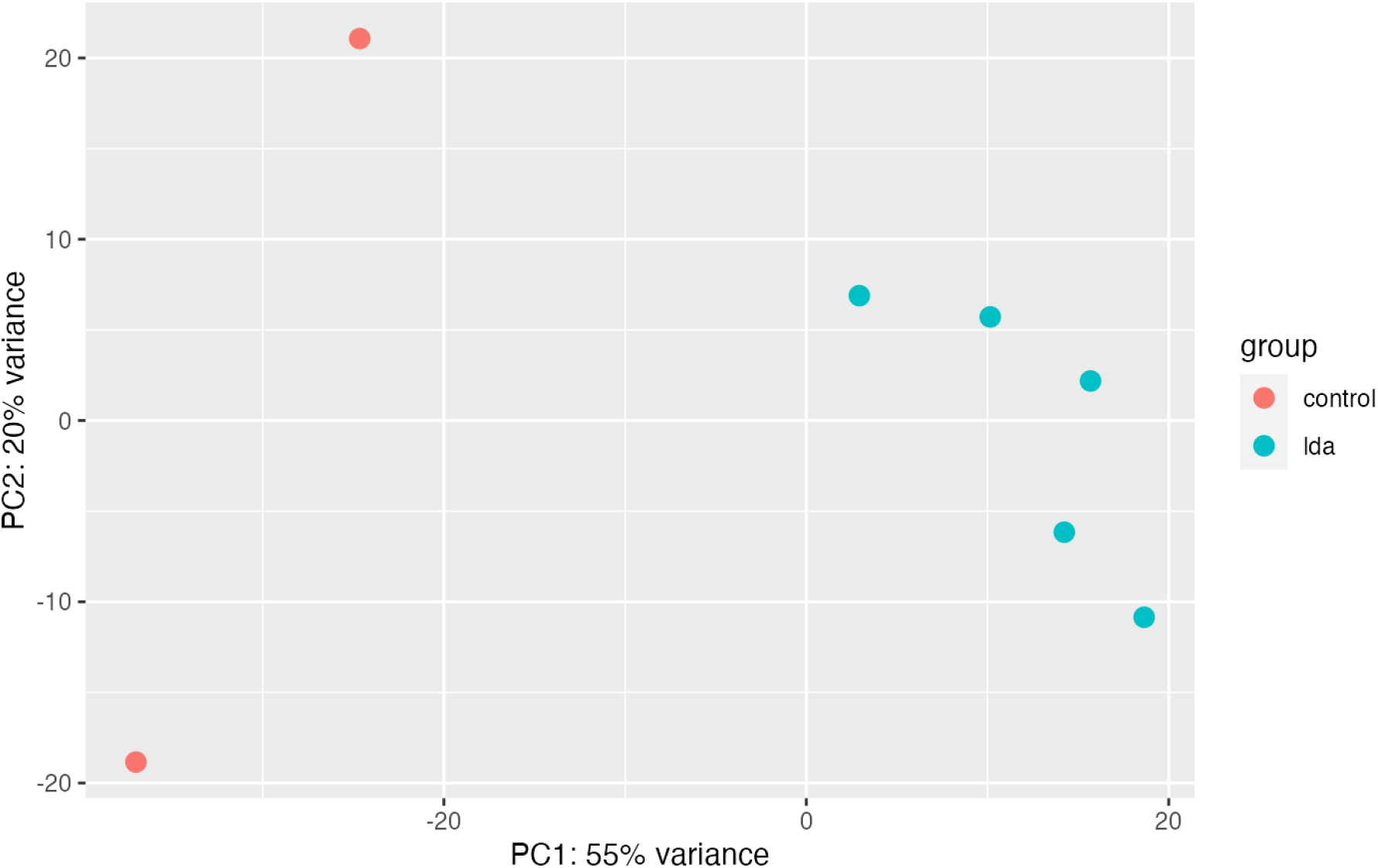
Principal component analysis (PCA) for control and LDA affected individuals. Principal components 1 and 2 were identified by variance stabilizing transformation in DESeq2.

The same clustering pattern can be observed with heatmap visualization of the expression profiles, so there is a well-defined distinction within the control and LDA affected groups. There is no grouping displayed when it comes to genotype or age (Figure 2).

**Figure 2.**
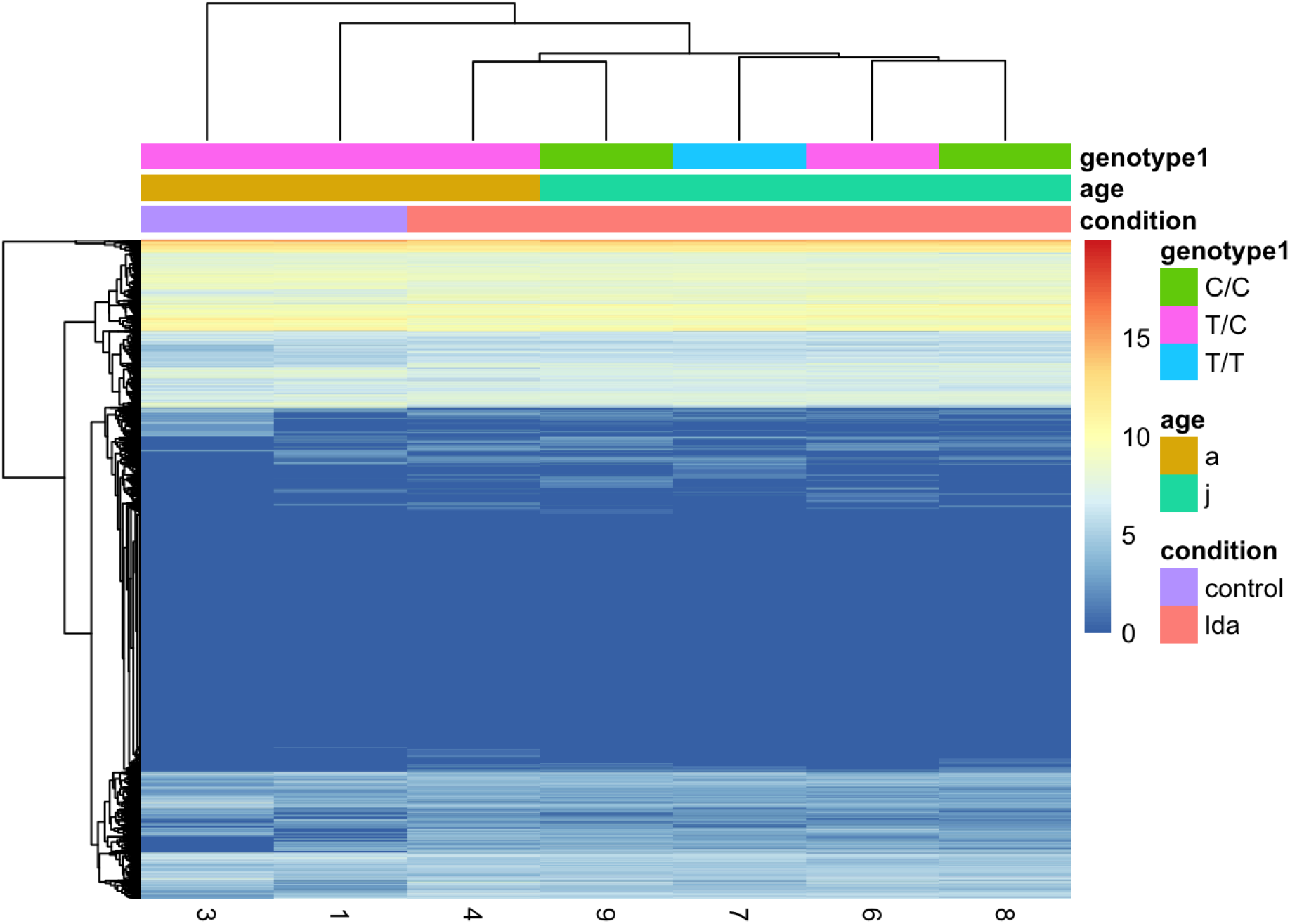
Heatmap analysis of expression profiles of control (1,3) and LDA affected animals. Legend displays genotypes (C/C, T/C and T/T), ages (adult, juvenile) and condition (control, LDA affected).

Volcano plot of differentially expressed genes reveals that there is a numerous difference between control and LDA affected animals in terms of expressed genes. The ten genes that have the highest LOg2FC values are UBL7, TTC33, PRKAA1, TXN2, PHOSPHO1, PSMG3, SNAPC4, LRRN4, NFIA and ADSL while the genes that had decreased expressions are JRK, FAM180B, ZNF660, MRPL33, SMAD5, RNF125, TP0, HSPB7, RAN, FAM200B (Figure 3.).

**Figure 3.**
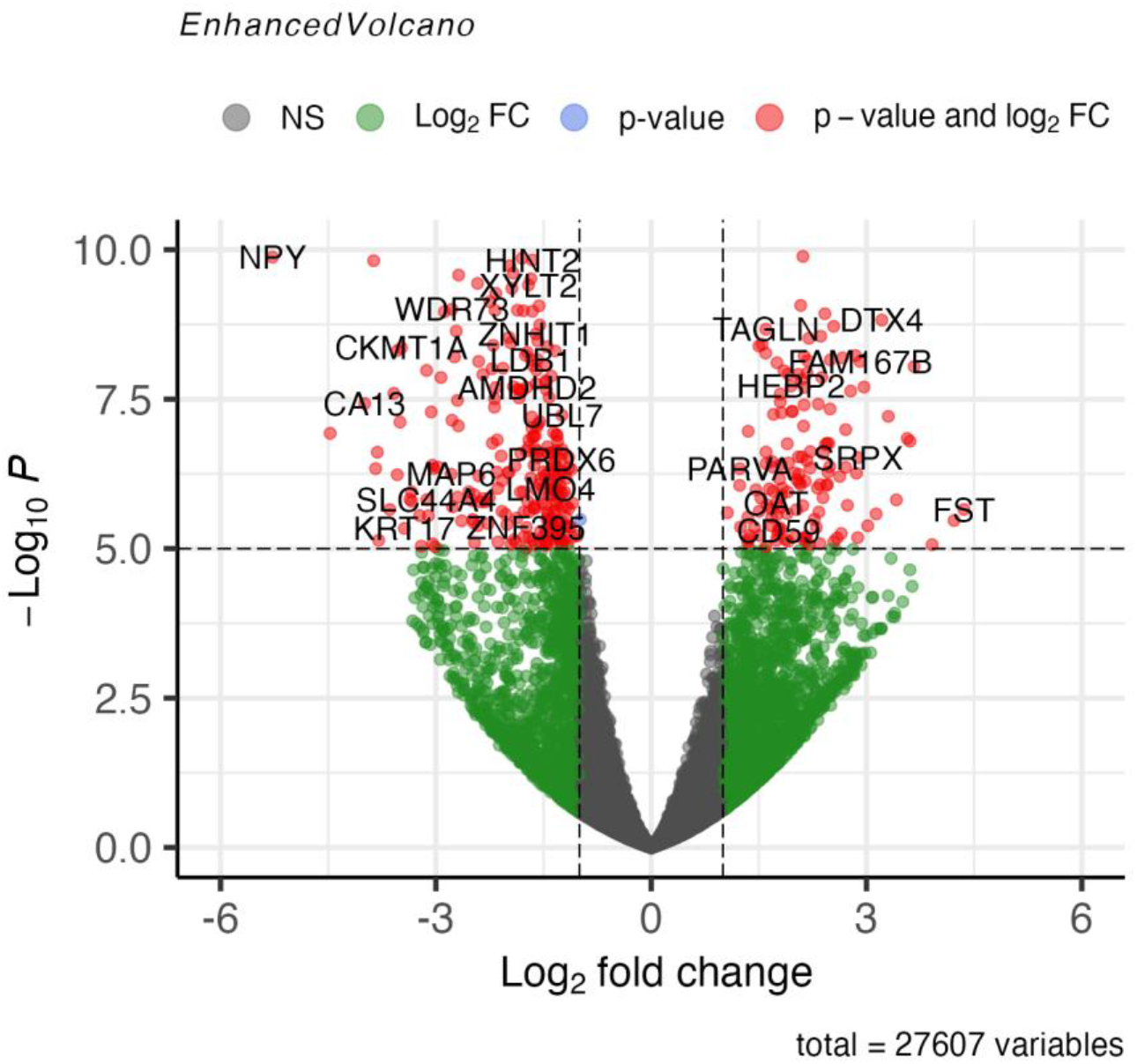
Volcano plot of differentially expressed genes. Genes with red are the ones that showed significantly different Log2FC.

Based on differentially expressed genes (DEGs) the Gene Ontology (GO) terms that were perturbed can be determined in a Gene Set Enrichment Analysis (GSEA) (Figure 4.). Overall, the majority of the suppressed GO terms (~40 genes in each) are involved in cellular respiration processes with p-values<0.0005. The activated GO terms are much more diverse in significance as well as in molecular function, biological processes.

**Figure 4.**
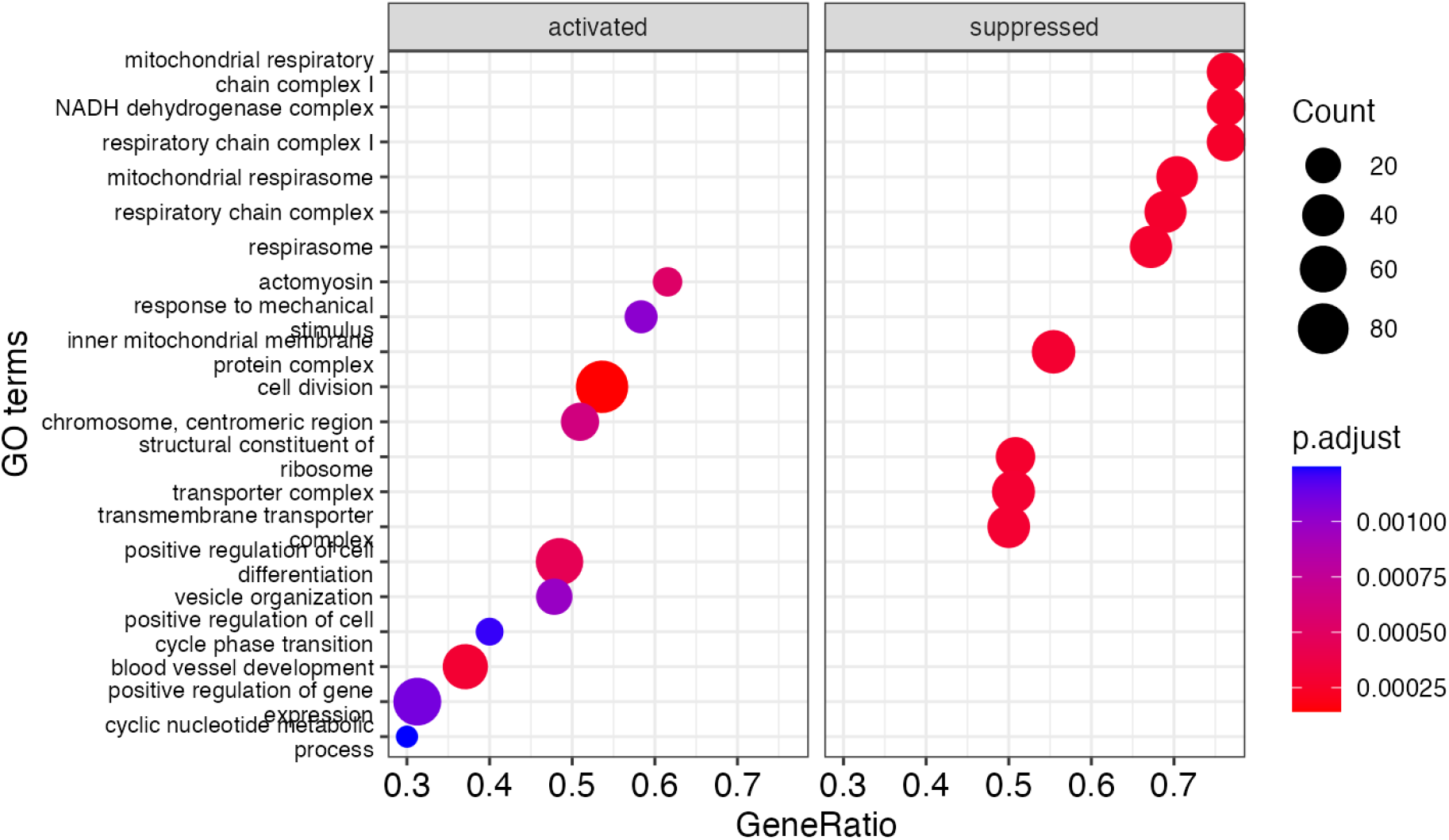
Gene Set Enrichment Analysis (GSEA) of the Gene Ontology terms based on differentially expressed genes.

The network visualization of GSEA shows that the nodes (GO terms) with higher connectivities are somehow related to cellular respiration processes (Figure 5.). These nodes are the same as the suppressed GO terms in the previous figure (Figure 4.).

**Figure 5.**
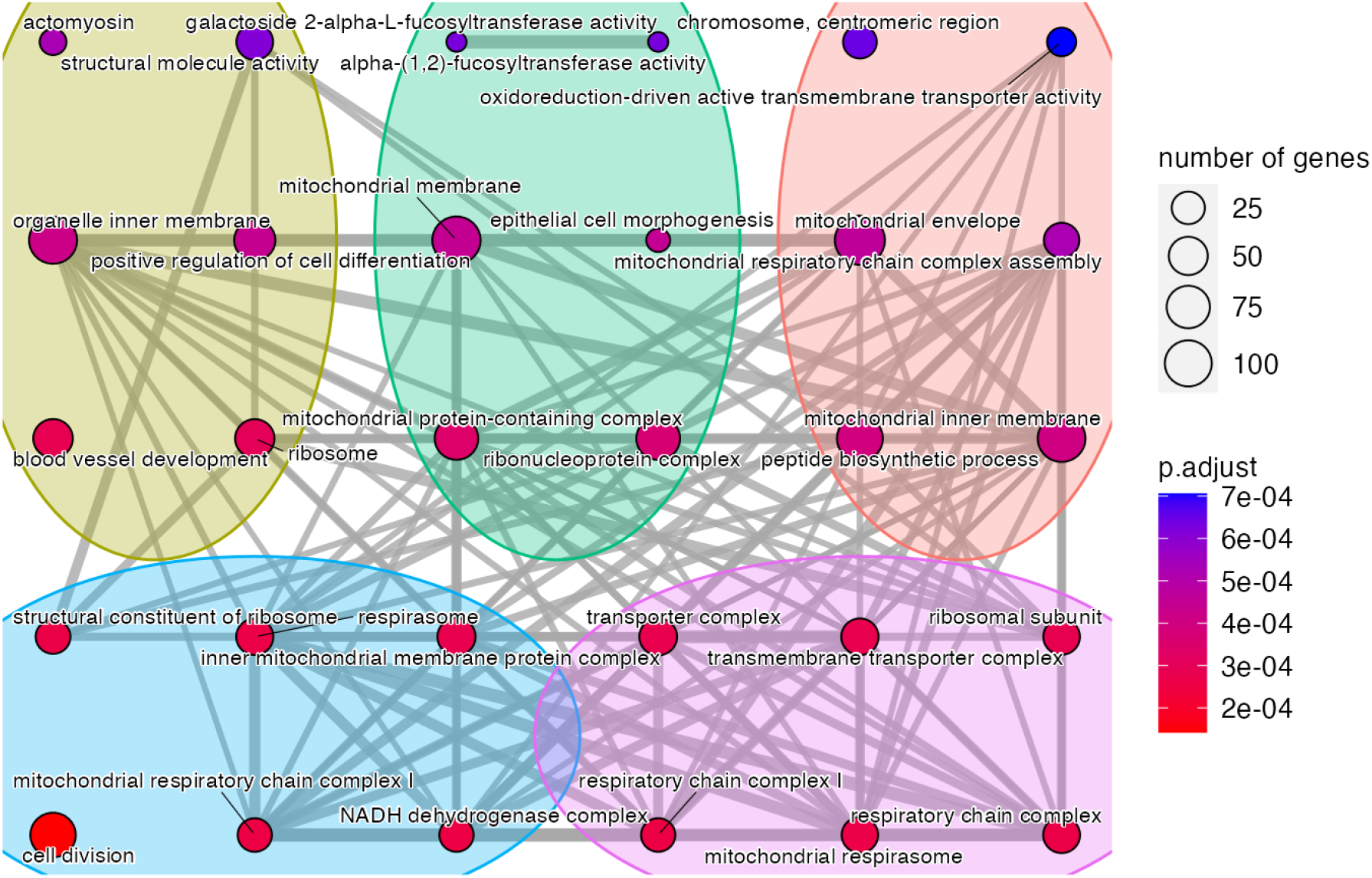
Network visualization of GSEA.

### Validation by RT-PCR and q-RT-PCR

For all the 9 gene (CAPN8, CCDC136, GKN2, GSDMC, ITGA5, ITGA7, MYL9, SMPD3, UGT1A1) we were able to verify the gene expression levels obtained during the RNA sequencing with q-RT-PCR (Figure 6.). The samples and all the primer pairs gave quantitatively identical expression compare to the RNA-Seq.

**Figure 6.**
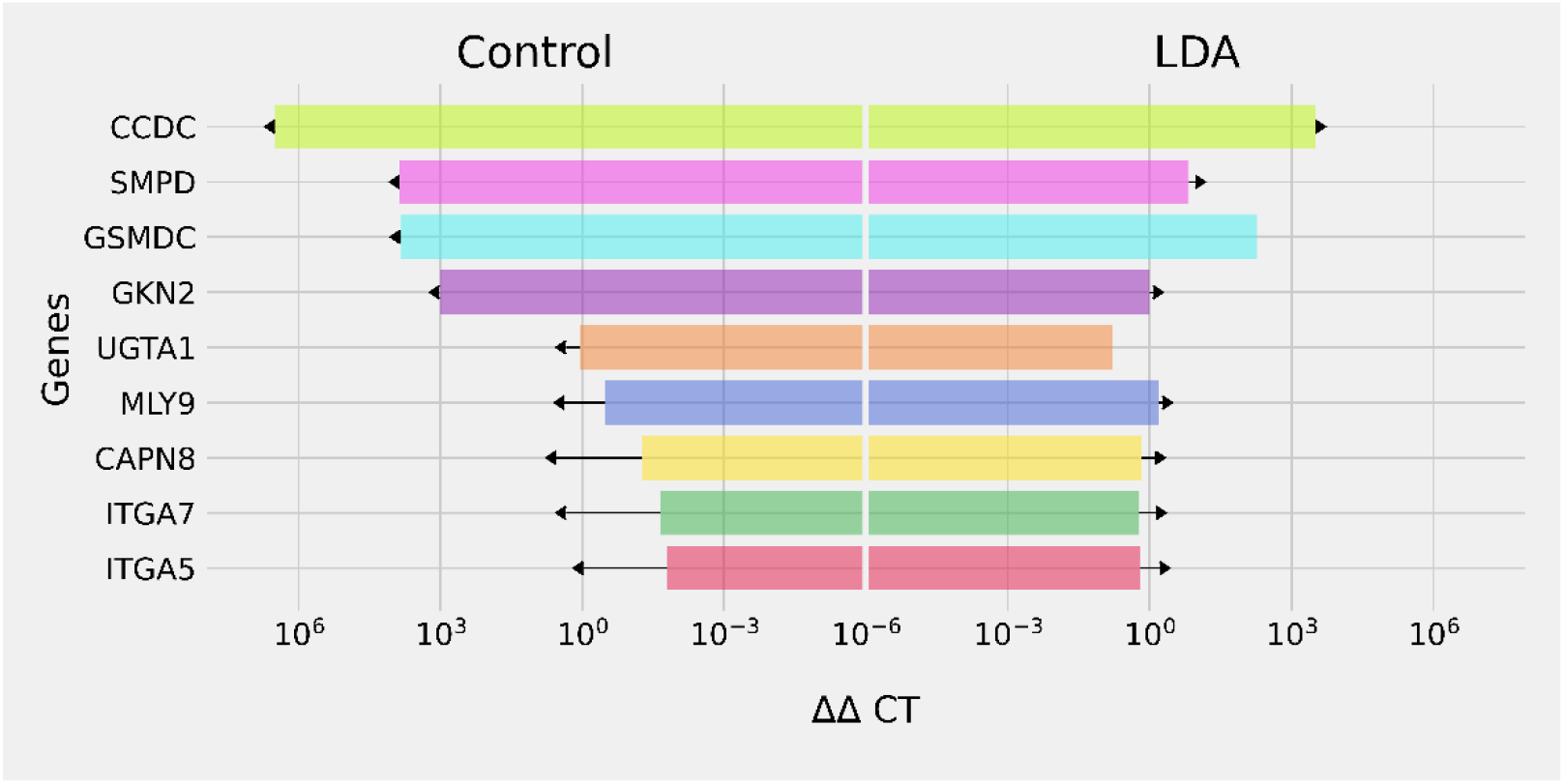
– q-RT-PCR data for 9 genes used as a validation of RNA sequencing.

Interestingly, the expression of the DOUX2 and DUOXA2 was reduced in the LDA affected animals compared to the control samples. These previous conclusions may serve as a basis for a better understanding of the diseases associated with hypomotility, but more precise characterization of the expression of DUAL oxidases is required in the cattle. Furthermore, it seems that in case of the disease the presence of these genes decreases highly.

## M & M

### Tissue sampling

All animal experimentation complied with the Hungarian Veterinary Authorities’ rules. We collected the samples in the dairy farm of Martontej Ltd, Ráczkeresztúr, Hungary. Abomasum tissue samples were collected from Holstein-Friesian cows during veterinary surgery, 5 of the animals was LDA diseased. After collecting the samples, we placed them to RNA Later solution and stored in -70 C until the RNA was isolated.

### RNA extraction and sequencing

Total RNA was extracted from all cattle tissues using RNAzol Reagent (RNazolRT Molecular Research Center) followed by DNase treatment and RNAqueous-Micro Kit RNA isolation kit (Invitrogen, Thermo Fisher Scientific, Waltham, USA) according to the manufacturer’s recommendation. The RNA integrity was verified using Agilent Bioanalyser 2100 (Agilent, Palo Alto, CA, USA). The samples were sequenced by UD-Genomed Ltd. using an Illumina NextSeq500 analyser system. We also extracted DNA from the abomasum tissue samples using phenol chloroform method for further genotyping works, like identify the genotype of the FN298674:g.90T>C SNP in MLN promoter by Sanger sequencing.

### Transcriptome assembly and gene expression

For quality control, FastQC (v0.11.9) was used to make sure that only samples with quality score above 28 were involved in further investigations. Bos taurus genome fasta (http://ftp.ensembl.org/pub/release-105/fasta/bos_taurus/dna/Bos_taurus.ARS-UCD1.2.dna.toplevel.fa.gz) and the corresponding GTF file (http://ftp.ensembl.org/pub/release-105/gtf/bos_taurus/Bos_taurus.ARS-UCD1.2.105.gtf.gz) were acquired from the FTP site of Ensembl genome browser. Modified version of Tuxedo protocol was used to carry out the initial steps of RNA sequencing pipeline which was partially proposed by Chu et al. (2017). This protocol uses HISAT2 (v2.2.1) to align reads to the given genome (Kim et al., 2019) and uses HTSeq (v0.11.3) to count the number of reads mapped to each genomic location (Putri et al., 2021). Gene count matrix was constructed with Python Pandas (v1.1.3). All the steps in the downstream analysis were performed in R. Differential gene expression analyses was done with DESeq2 (v1.34.0) (Love et al., 2014). For Gene Set Enrichment analysis Clusterprofiler (v4.2.2) was used (Wu et al., 2021). Ensembl id to external gene name transformation was carried out using biomaRt (v2.50.3) (Durinck et al., 2009).

### Validation by q-RT-PCR

We choose the genes ((DUOX2 (ENSBTAG00000016234), DUOXA2 (ENSBTAG00000016239), CAPN8 (ENSBTAG00000003600.6), CCDC136 (ENSBTAG00000011002.6), GKN2 (ENSBTAG00000017199.6), GSDMC (ENSBTAG00000017478.6), ITGA5 (ENSBTAG00000013745.6), ITGA7 (ENSBTAG00000012897.6), MYL9 (ENSBTAG00000011473.5), SMPD3 (ENSBTAG00000009639.5), UGT1A1 (ENSBTAG00000026181.5), RPS15A (ENSBTAG00000054615.1)) for validating the RNA sequencing data with q-RT-PCR. Expression level of these genes were significantly different between the healthy and the diseased group. Ribosomal protein s15 (RSP15A) was used as an endogenous control. All primer was designed with the Primer3Plus online designing tool on the basis of the cow sequences available from ensemble (RefSeq mRNA; Acc<u>:NM_001166500</u>) to obtained different length fragment from RNA as DNA, to check the DNA contamination of the samples. 200 ng RNA was reverse-transcribed to cDNA using High-Capacity cDNA Reverse Transcription kit (Applied Biosystems™, Thermo Fisher Scientific, Waltham, USA) in a 20 μL reaction including random primers, annealing and elongation incubations were conducted according to protocol. The q-RT-PCR reactions were conducted in 15 μL final volume including the 5x Hot FirePol EvaGreen qPCR Plus master mix (Solis BioDyne) and the reactions were run on Roche LifeCycler 96 System. All samples were tested in triplicated form. The DNA amplifications were performed with My Taq™ Red Mix (Meridian Bioscience, Cincinnati, USA). Initial denaturation (95°C for 1 min) was followed by 35 cycles of PCR: denaturation at 95°C for 20 sec; annealing at 58°C, 60°C for 20 sec, extension at 72°C for 20 sec. The final extension was 72°C for 2 minutes. The Table 1. contains the information of specific primers.

**Table 1.**
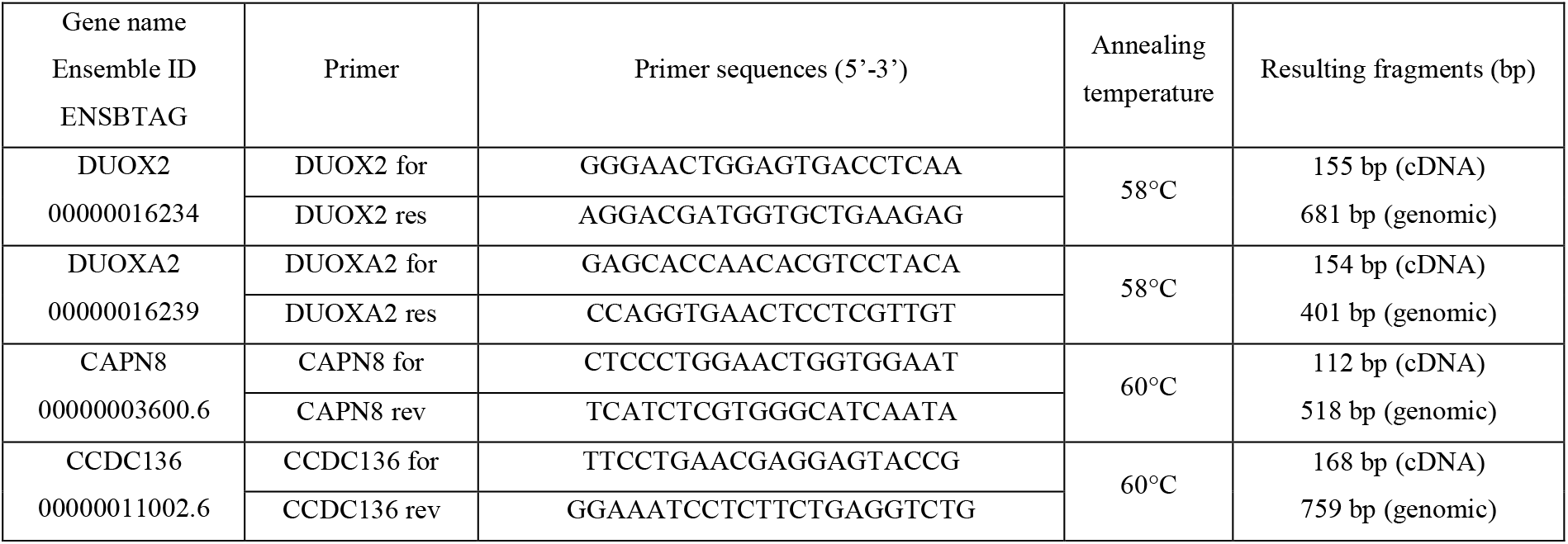
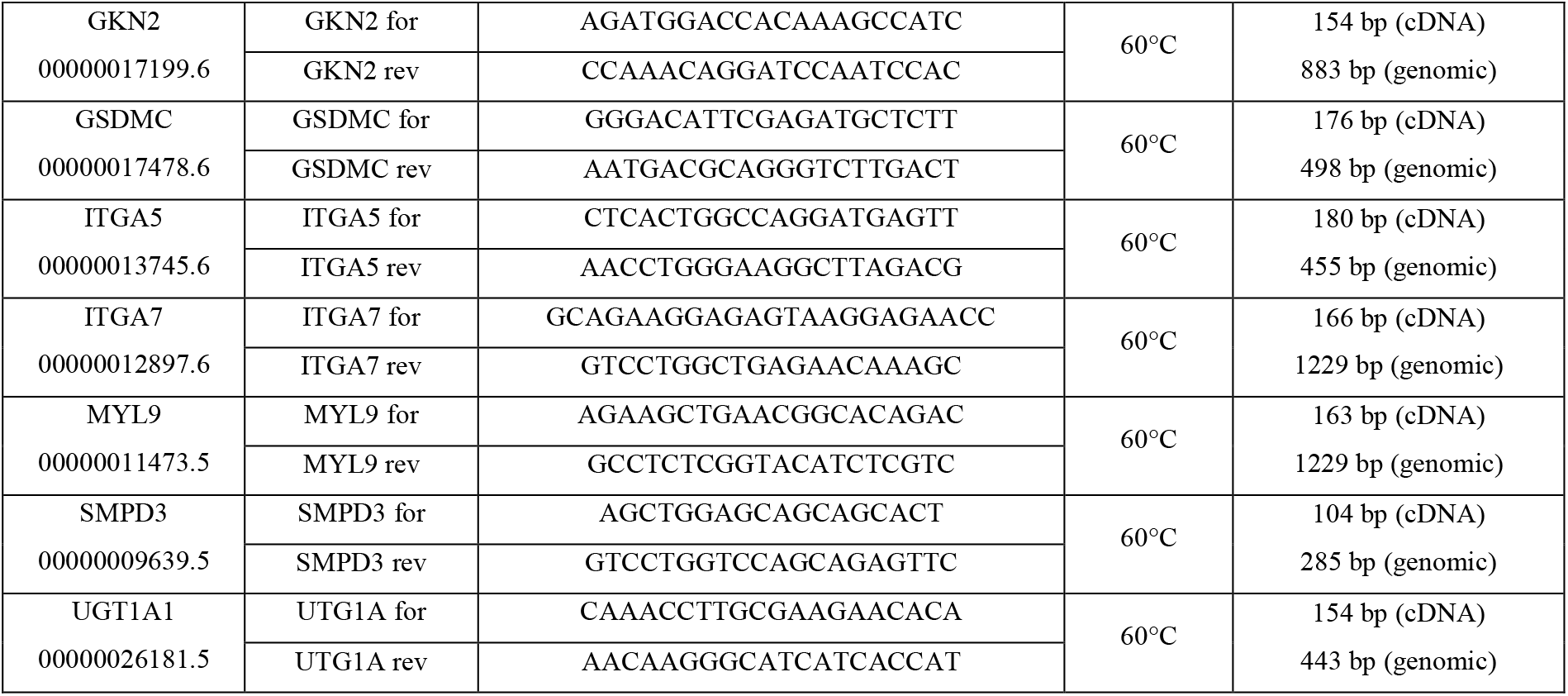
Oligonucleotide sequences

## Conclusion

Left-sided displacement of abomasum is a highly heritable disease affecting 1-7% of the herd of dairy farms, which is especially common in high-yielding animals (S.J. LeBlanc,2009). The incidence rate of the disease is increasing as we bring more and more productive individuals into breeding. OHV can occur at any age of the animal, most often it occurs around calving, when the movement of the animal’s intestinal system slows down, as a result of which gases accumulate in the abomasum, which moves out of its natural place. Sick animals produce up to 300 litres less milk during the lactation period, moreover, OHV requires surgical intervention, which greatly increases veterinary costs and the number of forced cullings (Song, 2020).

The present study demonstrates the transcriptome analysis of LDA diseased Holstein-Friesian animals. During the experiments performed with qRT-PCR and RNA sequencing, we were able to examine the level of DUOX2 expression in the abomasum of LDA-affected and healthy cattle. Interestingly, we found that the level of DUOX2 and DUOX2A gene expression was significantly reduced in animals suffering from LDA. Oxidative processes can lead to the formation of partially reduced, highly active oxygen metabolites, collectively known as reactive oxygen species (ROS). In mammalian species, superoxide and its derivatives are necessary to maintain proper homeostasis in the body, to moderate signal transduction, gene expression, and many other cellular processes. However, it is widely known that their excessive presence or contingent absence leads to the development of diverse diseases. As natural by-products of aerobic cellular metabolism, ROS maintain and regulate homeostasis and cell signalling. They are also essential for killing pathogens however, ROS can be harmful to host cells and tissues (Novo, 2008). Whether they act protective or detrimental, depends on the fine-tuning of the ROS production and disposal at the tissue, which is an evolutionarily conserved mechanism developed to be carefully balanced by generation and degradation of ROS. The ROS levels are strongly influenced by stress factors; the dysfunction of redox homeostasis results in increased production or inefficient elimination of ROS which causes oxidative stress. The sustained elevated level of oxidative stress can lead to significant cell damage causing apoptosis or inflammation and plays a significant role in the pathogenesis of chronic inflammation, activation of wound healing, and tissue fibrogenesis.

We examined the expression of the NOX family DUOX2 and its maturation factor DUOXA2. DUOX2/DUOX2A forms a stable complex on the cell surface and, like other NOX enzymes, participates in the formation of reactive oxygen radicals, more specifically, hydrogen peroxide. In addition to their role in redox processes, the ROS produced by DUOX enzymes support the immune system in the local fight against bacterial infections in the respiratory tract and lower tracts of the digestive system. DUOX enzymes play an essential role in the development of the immune response in the digestive system therefore loss-of-function mutations in the genes encoding the enzymes promote the progression of inflammatory bowel diseases in humans and in mouse models. The reduced functioning of the local immune response of the intestinal system can also cause or show the development of intestinal diseases in cattle, including LDA with a less clear background.

Our most important result is that we established that the expression level of the DUOX2 and DUOX2A genes belonging to the NOX gene family is significantly reduced in animals suffering from LDA. The experiments carried out add to the understanding of a disease that is a serious problem affecting the dairy farming sector. More information is needed in terms of both gene expression and microbiome for a more accurate understanding of the development and cause of the disease of bovine abomasum. Since the change in DUOX2/DUOX2A gene expression also leads to intestinal changes in other species and destroys the body’s resistance to bacterial infections, it may be a candidate gene for the development of the left displaced abomasum disease.

The next generation sequencing tools provide novel tools for expression profiling of biologically relevant networks and pathways. Here we identified the differentially expressed genes in the bovine abomasum in the presence of a disease whose genetic causes have not been precisely clarified yet.

## Acknowledgements

B.B. received NTP-NFTÖ-21-B-0127 scholarship. O.I.H. was funded by NKFIH OTKA FK 124708, NKFIH OTKA FK 111964, János Bolyai Research Scholarship of the Hungarian Academy of Sciences and New National Excellence Program of the Ministry for Innovation and Technology from the source of the National Research, Development and Innovation Fund (ÚNKP-21-5).

Thanks for Sándor Egedi, Zsuzsanna Galliné, Edit Gubó for the technical support.

## Authors’ contributions

RNA-Seq experiment and bioinformatic analysis carried out by BB, ZG and LK under the supervision of OIH. Validation of the bioinformatic analysis by qRT-PCR executed by ZSN ZG and BB. OIH conceived the study, analysed the data and drafted the manuscript. All authors read and approved the final manuscript.

## References

Attoub, S., Levasseur, S., Buyse, M., Goïot, H., Laigneau, J. P., Moizo, L., … & Bado, A. (1999). Physiological role of cholecystokinin B/gastrin receptor in leptin secretion. Endocrinology, 140(10), 4406–4410.

Brown, J. C., Johnson, L. P., & Magee, D. F. (1966). Effect of duodenal alkalinization on gastric motility. Gastroenterology, 50(3), 333–339.

Brown, J. C., Mutt, V., & Dryburgh, J. R. (1971). The further purification of motilin, a gastric motor activity stimulating polypeptide from the mucosa of the small intestine of hogs. Canadian journal of physiology and pharmacology, 49(5), 399–405.

Chu, C. P., Hokamp, J. A., Cianciolo, R. E., Dabney, A. R., Brinkmeyer-Langford, C., Lees, G. E., & Nabity, M. B. (2017). RNA-seq of serial kidney biopsies obtained during

Darvodelsky, A. M., Davey, M. W., Reid, A. M., Titchen, D. A., & Wang, X. (1988). Immunochemical characterisation of somatostatin in ruminants. Regulatory peptides, 20(2), 161–170.

Deloose, E., Verbeure, W., Depoortere, I., & Tack, J. (2019). Motilin: from gastric motility stimulation to hunger signalling. Nature Reviews Endocrinology, 1.

Durinck, S., Spellman, P. T., Birney, E., & Huber, W. (2009). Mapping identifiers for the integration of genomic datasets with the R/Bioconductor package biomaRt. Nature protocols, 4(8), 1184–1191.

Egloff, P., Hillenbrand, M., Klenk, C., Batyuk, A., Heine, P., Balada, S., … & Plückthun, A. (2014). Structure of signaling-competent neurotensin receptor 1 obtained by directed evolution in Escherichia coli. Proceedings of the National Academy of Sciences, 111(6), E655–E662.

Hamann, H., Wolf, V., Scholz, H., & Distl, O. (2004). Relationships between lactational incidence of displaced abomasum and milk production traits in German Holstein cows. Journal of Veterinary Medicine Series A, 51(4), 203–208.

Hauge-Evans, A. C., King, A. J., Carmignac, D., Richardson, C. C., Robinson, I. C., Low, M. J., … & Jones, P. M. (2009). Somatostatin secreted by islet δ-cells fulfills multiple roles as a paracrine regulator of islet function. Diabetes, 58(2), 403–411.

Huang, Z. G., Xiong, L., Zhen-Shan, L. I. U., Yong, Q. I. A. O., Rong, D. A. I., Zhuang, X. E., … & Guo-Qing, L. I. U. (2006). The tissue distribution and developmental changes of ghrelin mRNA expression in sheep. Acta Genetica Sinica, 33(9), 808–813.

Jonnakuty, C., & Gragnoli, C. (2008). What do we know about serotonin?. Journal of cellular physiology, 217(2), 301–306.

Keskin, O., & Yalcin, S. (2013). A review of the use of somatostatin analogs in oncology. OncoTargets and therapy, 6, 471.

Kim, D., Paggi, J. M., Park, C., Bennett, C., & Salzberg, S. L. (2019). Graph-based genome alignment and genotyping with HISAT2 and HISAT-genotype. Nature biotechnology, 37(8), 907-915.progression of chronic kidney disease from dogs with X-linked hereditary nephropathy. Scientific reports, 7(1), 1–14.

Novo, E., Parola, M. Redox mechanisms in hepatic chronic wound healing and fibrogenesis. Fibrogenesis Tissue Repair 1, 5 (2008). https://doi.org/10.1186/1755-1536-1-5

Leiter, A. B., Keutmann, H. T., & Goodman, R. H. (1984). Structure of a precursor to human pancreatic polypeptide. Journal of Biological Chemistry, 259(23), 14702–14705.

Love, M. I., Huber, W., & Anders, S. (2014). Moderated estimation of fold change and dispersion for RNA-seq data with DESeq2. Genome biology, 15(12), 1–21.

Mömke, S., Sickinger, M., Rehage, J., Doll, K., & Distl, O. (2012). Transcription factor binding site polymorphism in the motilin gene associated with left-sided displacement of the abomasum in German Holstein cattle. PloS one, 7(4), e35562.

Mömke, S., Sickinger, M., Lichtner, P., Doll, K., Rehage, J., & Distl, O. (2013). Genome-wide association analysis identifies loci for left-sided displacement of the abomasum in German Holstein cattle. Journal of dairy science, 96(6), 3959–3964.

Putri, G. H., Anders, S., Pyl, P. T., Pimanda, J. E., & Zanini, F. (2021). Analysing high-throughput sequencing data in Python with HTSeq 2.0. arXiv preprint 2112.00939.

Ratner, C., Hundahl, C., & Holst, B. (2016). The metabolic actions of neurotensin secreted from the gut. Cardiovascular Endocrinology, 5(3), 102–111.

Reynolds, G. W., Hansky, J., & Titchen, D. A. (1984). Immunoreactive gastrin concentrations in gastrointestinal tissues of sheep. Research in veterinary science, 37(2), 172–174.

Romański, K. W. (2017). Importance of the enteric nervous system in the control of the migrating motility complex. Physiology international, 104(2), 97–129.

S.J. LeBlanc, K.E. Leslie, T.F. Duffield, Metabolic Predictors of Displaced Abomasum in Dairy Cattle, Journal of Dairy Science, Volume 88, Issue 1, 2005, Pages 159–170, ISSN 0022-0302.

Song, Y., Loor, J.J., Zhao, C. et al. Potential hemo-biological identification markers to the left displaced abomasum in dairy cows. BMC Vet Res 16, 470 (2020). https://doi.org/10.1186/s12917-020-02676-x

Trapnell C, Pachter L, Salzberg SL: TopHat: discovering splice junctions with RNA-Seq. Bioinformatics. 2009, 25 (9): 1105–1111

Trapnell C, Williams BA, Pertea G, Mortazavi A, Kwan G, Van Baren MJ, Salzberg SL, Wold BJ, Pachter L: Transcript assembly and quantification by RNA-Seq reveals unannotated transcripts and isoform switching during cell differentiation. Nat Biotechnol. 2010, 28 (5): 511–515.

Van Winden, S. C., & Kuiper, R. (2003). Left displacement of the abomasum in dairy cattle: recent developments in epidemiological and etiological aspects. Veterinary research, 34(1), 47–56.

Wang, W., Cheng, L., Guo, J., Ma, Y., & Li, F. (2014). Expression of Ghrelin in gastrointestinal tract and the effect of early weaning on Ghrelin expression in lambs. Molecular biology reports, 41(2), 909–914.

Wolf, V., Hamann, H., Scholz, H., & Distl, O. (2001). Influences on the occurrence of abomasal displacements in German Holstein cows. DTW. Deutsche tierarztliche Wochenschrift, 108(10), 403–408.

Wu, T., Hu, E., Xu, S., Chen, M., Guo, P., Dai, Z., … & Yu, G. (2021). clusterProfiler 4.0: A universal enrichment tool for interpreting omics data. The Innovation, 2(3), 100141.

